# Generation of *de novo* miRNAs from template switching during DNA replication

**DOI:** 10.1101/2023.04.19.537475

**Authors:** Heli A. M. Mönttinen, Mikko J. Frilander, Ari Löytynoja

## Abstract

The mechanisms generating novel genes and genetic information are poorly known, even for microRNA (miRNA) genes with an extremely constrained design. All miRNA primary transcripts need to fold into a stem-loop structure to yield short gene products (∼22 nt) that bind and reppress their mRNA targets. While a substantial number of miRNA genes are ancient and highly conserved, short secondary structures coding for entirely novel miRNA genes have been shown to emerge in a lineage-specific manner. Template switching is a DNA-replication-related mutation mechanism that can introduce complex changes and generate perfect base pairing for entire hairpin structures in a single event. Here we show that the template-switching mutations (TSMs) have participated in the emergence of over 6,000 suitable hairpin structures in the primate lineage to yield at least 18 new human miRNA genes, that is 26% of the miRNAs inferred to have arisen since the origin of primates. While the mechanism appears random, the TSM-generated miRNAs are enriched in introns where they can be expressed with their host genes. The high frequency of TSM events provides raw material for evolution. Being orders of magnitude faster than other mechanisms proposed for *de novo* creation of genes, TSM-generated miRNAs enable near-instant rewiring of genetic information and rapid adaptation to changing environments.

---

The emergence of novel genes from already pre-existing ones during evolution is a rather common and well-characterized process involving a variety of mechanisms such as gene fusions and fissions, horizontal gene transfers, gene duplication and insertions of mobile DNA elements (1, 2). However, the *de novo* emergence of new genes has been deemed rare, particularly with protein-coding genes that need to satisfy the requirement for both transcription and translation processes and post-transcriptional RNA processing events such as pre-mRNA splicing and polyadenylation (3, 4). Non-coding RNA (ncRNA) genes have no translational requirement and are thus more relaxed to emerge and evolve (5, 6). Among the ncRNAs, genes encoding microRNAs (miRNAs) are unique in gene structure and function (7). The primary transcript from a miRNA locus folds to an RNA stem-loop structure with a strong preference for 35 bp stem length and 10 nt or longer unstructured apical loop (8). It then undergoes both nuclear and cytoplasmic processing events that result in the final ∼22-nt miRNA gene product loaded on the RNA-induced silencing complex (8, 9). Ultimately, the functionality of a miRNA is determined by the potential for base-pairing interactions with an mRNA. In animals, miRNAs associate predominantly with the 3’UTR sequence of an mRNA through partial base-pairing interactions. Due to the flexibility of the mRNA recognition, a single miRNA can target multiple mRNAs (7).

A substantial number of miRNA genes are ancient and highly conserved, but many are subject to rapid evolutionary change (10) and novel miRNA genes have been shown to emerge in a lineage-specific manner e.g. in primates (11) and plants (12). The *de novo* formation of a miRNA gene requires a mutational mechanism that can introduce inverted DNA sequences to a transcribed region of the genome. Recent studies on mutation clusters within primate genomes have revealed the emergence of short inverted DNA sequences through a DNA replication-related mutation mechanism (13–15). This mechanism was initially proposed to explain the conversion of quasi-palindromic sequences into perfect palindromes by template-strand switching during replication (16, 17). The outcomes from such template switching are a loopback hairpin formation in the growing strand (intra-strand switch) or a brief jump to the opposing strand (inter-strand switch), followed by continuation of the DNA replication on the original strand (Fig. 1A). In both cases the result is a one-step formation of a complex replacement mutation, creating either an inverted repeat, an inversion, or a combination of inverted and direct repeats (Fig. S1). Hereafter, we refer to this as a template-switching mutation (TSM).

**Fig. 1.**
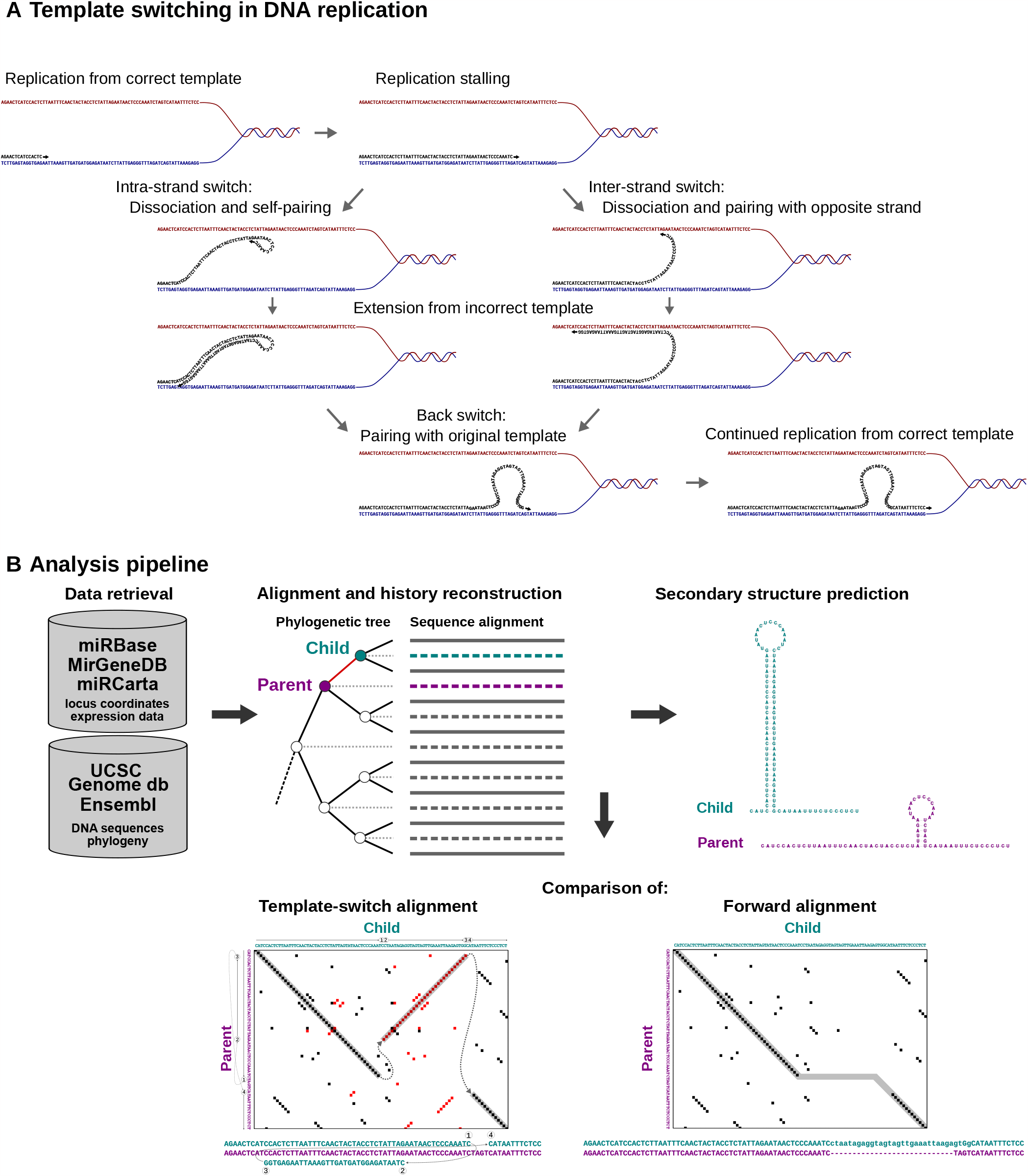
Mutation mechanism and analysis workflow. (*A*) TSMs are thought to result from DNA replication bypassing DNA lesions with a jump to the other DNA strand. After the replication stalling, the nascent strand may either fold over and undergo intramolecular pairing (Intra-strand switch) or pair with the opposite strand (Inter-strand switch); the nascent strand is then briefly extended from the incorrect template before realigning with the original template to continue the replication normally. In addition to a complex sequence change (shown as a loop), this may create a reverse-complement repeat with the potential to form a perfect hairpin. (*B*) For each miRNA in miRBase, the surrounding genome sequences were fetched for a set of mammals, and multiple alignment and sequence history (dashed lines in the sequence alignment) were inferred following a consensus phylogeny. The resulting tree-sequence structure was traversed and each parent-child sequence pair (here shown for the branch indicated with red) was searched for complex mutations. Mutation loci where the template-switch alignment was superior to the traditional forward alignment were recorded (bottom) and secondary structures before and after the inferred TSM event were predicted (right). The example is based on hsa-mir-5690.

We hypothesized that, in suitable circumstances, the TSM-generated inverted repeats could evolve into miRNA genes. To test this hypothesis, we reconstructed detailed evolutionary histories for the annotated human miRNA genes and studied the mutation steps ultimately resulting in novel, experimentally validated genes. We then assessed the dynamics of these emerging genes and the step leading to their regulation and biological function to understand how random mutations create biological complexity and new genetic information.

## Results

### Template switch mutations create and modify miRNA stem structures

We studied the emergence of miRNA genes by reconstructing evolutionary histories for the gene loci across mammals, focusing on the human lineage and the densely sampled primates. Applying the approach described earlier (18), we reconstructed the sequence history of all 1,821 unique human miRNA loci present in miRBase (v. 22.1; 19) and the 567 loci passing the more stringent annotation of MirGeneDB (v. 2.1; 20, 21). Given the phylogenetic tree-sequence alignment structure (Fig. 1B), we systematically traversed the tree and compared pairwise each Parent-Child pair. More specifically, we assumed that the ‘Parent’ sequence had evolved to ‘X’ which was converted by a TSM to ‘Y’, and this then evolved into the ‘Child’ sequence, in short: Parent 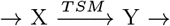 Child. As ‘X’ and ‘Y’ cannot be observed, we used the inferred ‘Parent’ and ‘Child’ sequences as their proxies. We found 1,633 of the miRBAse miRNA loci to have either emerged or significantly changed (>10% sequence change, see Methods) in the primate lineage. We then tested whether TSM could provide a more parsimonious explanation for the complex differences observed between the sequences at a parental and child node rather than traditional point substitutions and insertion-deletions (indels; see Fig. 1B). We considered TSM solutions that included an inverted fragment of ≥15 nucleotides and explained ≥5 sequence mismatches (see Methods). As ancestral reconstruction loses accuracy with high sequence divergence (22), we applied strict criteria for the implied sequence homology and conservation.

Among the miRBase cases passing our criteria, we found 79 TSMs that fully and 30 TSMs that partially (≥ 10 nucleotides are left unexplained by the primary TSM; see Methods) explain the *de novo* creation of miRNA stem structure at a given locus. Additionally, we found four cases where TSM had modified an existing miRNA hairpin by inverting its loop sequence, two of these at loci with a hairpin created by an earlier TSM (Table S1, Table S2, Data S1). Hsa-mir-4713 is an example that combines both types of events (Fig. 2A,B): the initial emergence of a miRNA stem in the ancestor of primates followed by an inversion of the loop sequence in the ancestor of apes. The first event is explained by a 28-base-long TSM (Fig. 2C), which may have occurred either intra- or intermolecularly (see Fig. 1A). In contrast, the second event is consistent with an intermolecular TSM of up to 83 bases (Fig. 2C). Of the 111 TSM-explained miRBAse loci, 18 were included in the more stringently annotated MirGeneDB (Table S1, Data S1).

**Fig. 2.**
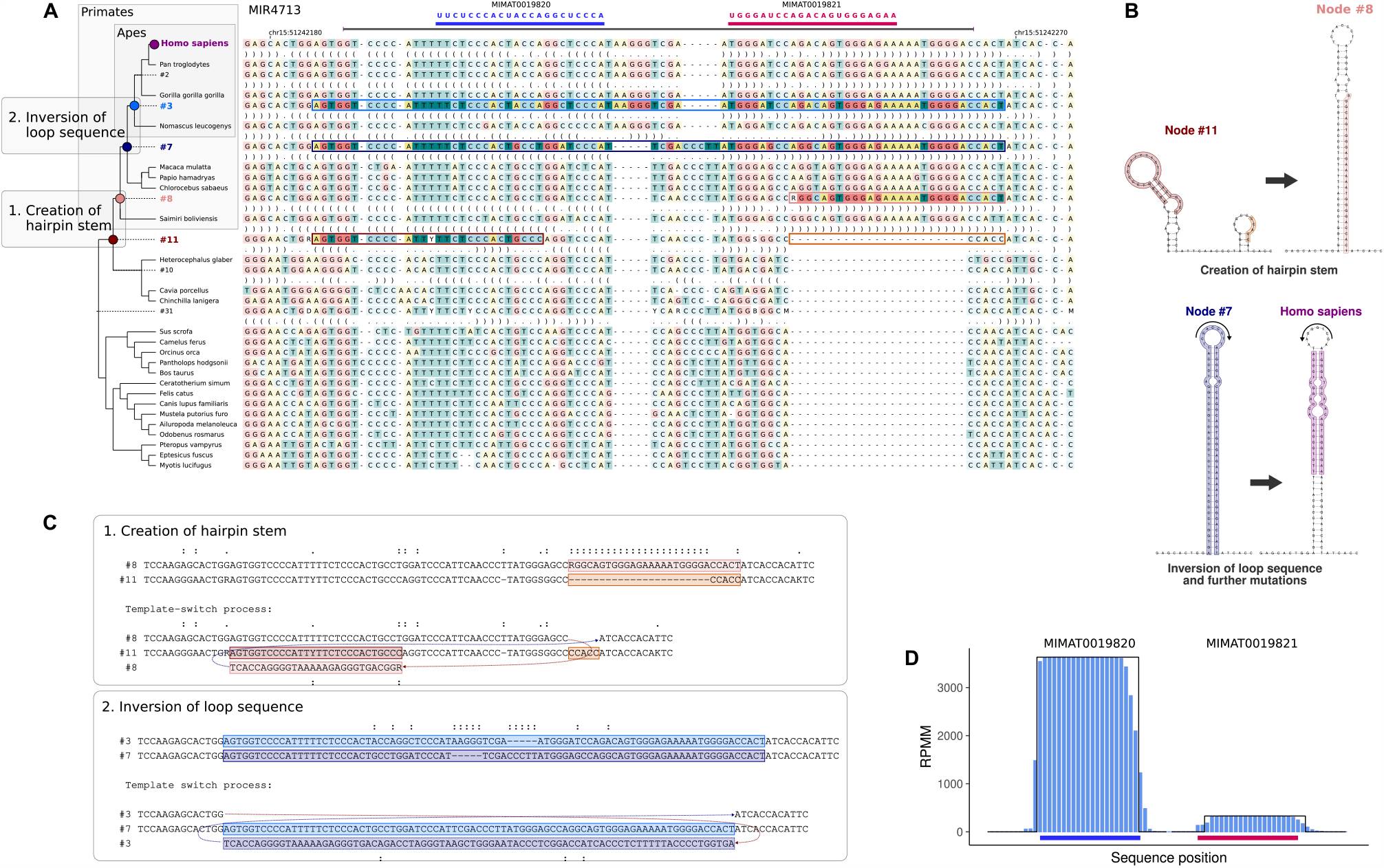
Emergence and evolution of the hsa-mir-4713 gene. (*A*) Two TSM events are inferred to have taken place in the locus among the sampled mammals. The first event in the ancestor of primates (between nodes #11 and #8) creates a new inverted repeat with the capacity of forming a hairpin. The second event at the root of apes (between nodes #7 and #3) modifies the existing hairpin by inverting the hairpin loop sequence. (*B*) Predicted secondary structures for the sequences before and after the inferred TSM events. Mismatches in the predicted stems are caused by inaccuracies in the sequence reconstruction and independent point mutations in-between the TSM event and the next internal node in the tree. (*C*) The inferred template switch solutions for the two complex mutations. The sequence names (left) match the node names in the tree. (*D*) Expression support for the hsa-mir-4713 gene in the miRCarta database. Light blue bars show the read counts (y-axis) for the two annotated transcripts (x-axis; blue and red bars, see panel *A*. Black lines indicate the ideal stacks with uniform counts (either zero or the maximum count) used for scoring the locus.

### Stable miRNA stem structures emerge from unstructured sequences

Although template switching was originally proposed as a mechanism for converting nearly perfect reverse-complement repeats into perfect ones (16, 17, 23, 24), it has been shown to take place also at loci with no pre-existing similarity (13–15). To investigate the requirement for pre-existing repeat sequences, we assessed the genomic context and the impact of the inferred TSMs by predicting secondary structures for the loci before and after the inferred events (Fig. 1B, ‘Secondary structure prediction’). Of the 1,366 primate-evolved miRNA loci, we focused on the 79 where template switching fully explains the emergence of the stem structure (Table S1, Table S2) and identified homologous sequences in a set of mammalian genomes. For these, we predicted the RNA secondary structures and determined the corresponding lowest free energy values.

The inferred TSM events are associated with a sharp drop in free energy, indicating a sudden appearance of complex secondary structures (Fig. 3). All of the inferred events occurred within the primate subtree: 57/79 took place in the ancestor of all primates; 11 and 5 in the ancestor of Old World monkeys and apes, respectively; and the last 6 stem structures are only present in a subset of apes (Fig. 3, Table S2). Of these miRNAs, 13 are included in the more strictly defined set of miRNA genes in MirGeneDB. Consistent with our phylogenetic analysis, MirGeneDB annotates all 13 miRNAs to have emerged either in Old World monkeys or human (Fig. 3) and lists each as the only member of a miRNA family of its own, underlining their uniqueness and *de novo* origin. Recently, the phylogenetic origin of MirGeneDB genes has been inferred in greater detail using covariance model-based methods (25, 26). For 12 of the 13 TSM-loci, our phylogenetic classification is consistent with that of ncOrtho (25). The exception is hsa-mir-4731 that has clearly arisen by a TSM in the ancestor of Homininae (Data S1) but is inferred by ncOrtho to originate in early primates. The erroneous inference may be explained by the conservation of the 5’ part of the gene and the potential for the locus to form some secondary structure before the TSM event (Data S1).

**Fig. 3.**
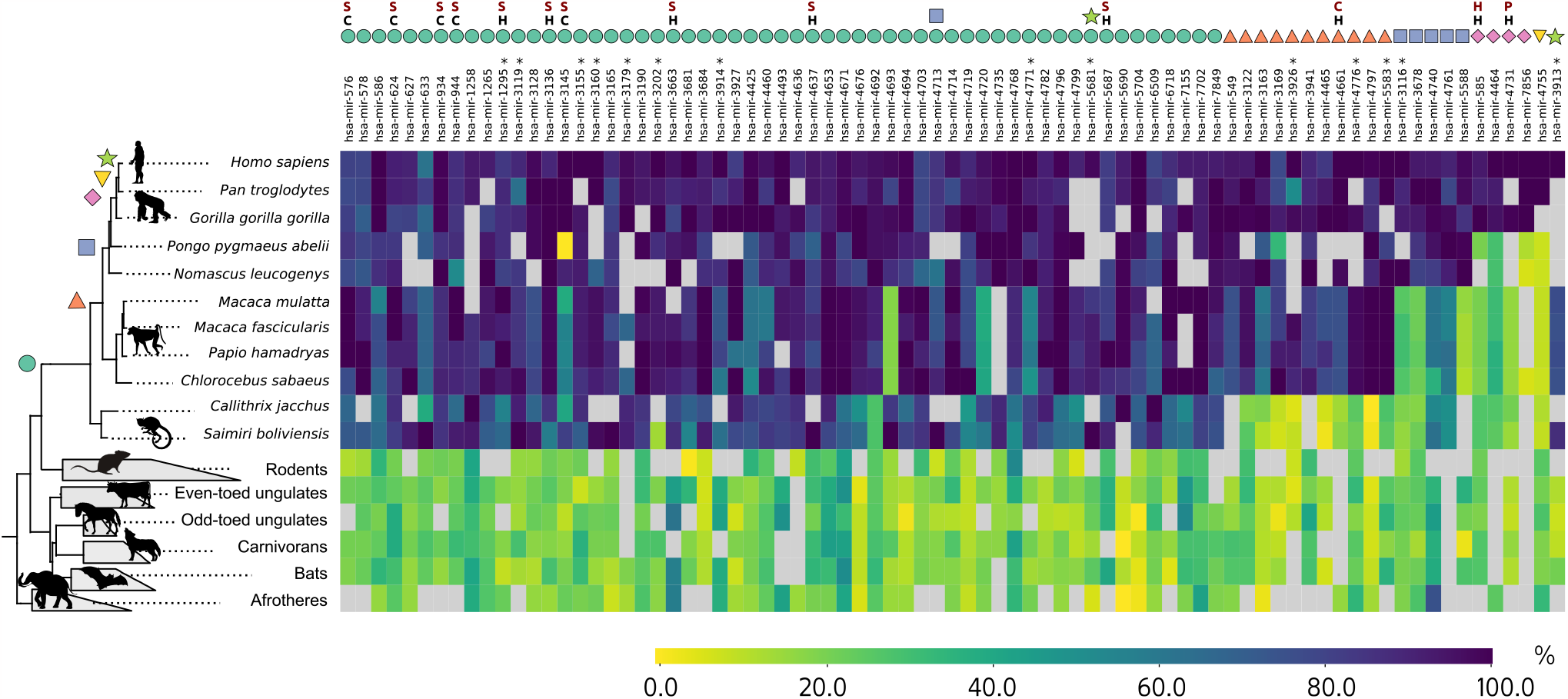
Normalised free-energy change at human miRNA loci. Matrix cells show the relative stability (free-energy change) of the predicted secondary structures for sequences (species; y-axis) homologous to the human miRNA loci (x-axis, top) in comparison to the sequence with the greatest free-energy change. Darker colours indicate greater changes and thermodynamically more stable secondary structures. Taxa with no homologous sequence are shown in grey. Outside of primates, species within the order are merged (grey trapezoids) and, within the merged group, the species with the greatest free-energy change is shown. Only taxa with data for at least 10 loci are included. Coloured symbols on the top indicate the tree branch where the TSM event was inferred to have taken place, in two cases the mutation hitting the same locus twice. Genes found in MirGeneDB are marked with black C and H, indicating that MirGeneDB reports their node of origin to be either Catarrhini (Old World monkeys) or *H. sapiens* (human), and with red P, S, C and H, indicating that ncOrtho infers their node of origin to be either Primates, Simiiformes (higher primates), Catarrhini (Old World monkeys), or Homininae (African apes). Asterisks after the gene names indicate loci with multiple annotated miRNA transcripts. See Fig. S2 for the full matrix.

On the other way around, MirGeneDB and ncOrtho list 83 and 69, respectively, of the 266 human miRNA families to have originated since the split of primates. We infer 18 of these novel primate-specific miRNA genes to have been generated by TSMs. This number may be an underestimate, however, as many primate-specific miRNA genes are located in genomic regions for which no homology is identified outside the primates and thus no ancestral reconstruction and inference of the mutation process can be performed. Moreover, we detected cases that can be explained by multiple template switches (Fig. S3) and, by requiring more than the two reciprocal jumps between the templates, are not captured by our current algorithm. Such complex mutations are not unprecedented and have been reported behind human genetic disorders (27).

### TSM-generated miRNAs are found in other evolutionary lineages

With the current species set and the quality control approach, we should theoretically be able to find the TSM events all the way to the ancestor of primates and rodents. However, our identification of novel miRNAs in primates only does not need to imply that the mechanism is primate-specific but more likely reflects the limitations of the underlying data. More specifically, ancestral sequence reconstruction inevitably loses accuracy in deeper nodes (22) and the exceptionally fast evolution of rodent sequences (28) makes identification of homologues difficult (cf. Fig. 3, grey cells). To assess whether the generation of miRNAs by the TSM mechanism takes also place outside primates, we repeated parts of the analyses for an alternative set of densely sampled full genomes, the laboratory mice and their relatives.

The Murinae set is phylogenetically more limited than the primates set and contains 16 strains of *M. musculus*, four other species from the genus *Mus*, and the rat as an outgroup. We replicated the search for TSM-generated stem structures at the 1,132 mouse miRNA loci annotated in miRBase. Although we could only identify cases that have taken place in the genus *Mus*, we found ten strong candidates for TSM-generated miRNA loci, four of which are annotated in MirGeneDB (Data S2). Mmu-mir-3086 is an example of a young miRNA present in a subset of species (Fig. 4A): The gene is listed both in miRBase and MirGeneDB and is inferred to have arisen by two independent TSM events, the first generating the hairpin structure and the second inverting the sequence in place (Fig. 4B,C). The miRNA is reported to show robust Drosha-dependent processing (29), and is predominantly expressed particularly in the mouse brain and embryonic stem cells (29). The regions coding for the miRNA products are perfectly conserved across the four *Mus* species with the exception of a double mutation in the 5’ product of the lab strain PWK/PhJ (Fig. 4A, red dashed box).

**Fig. 4.**
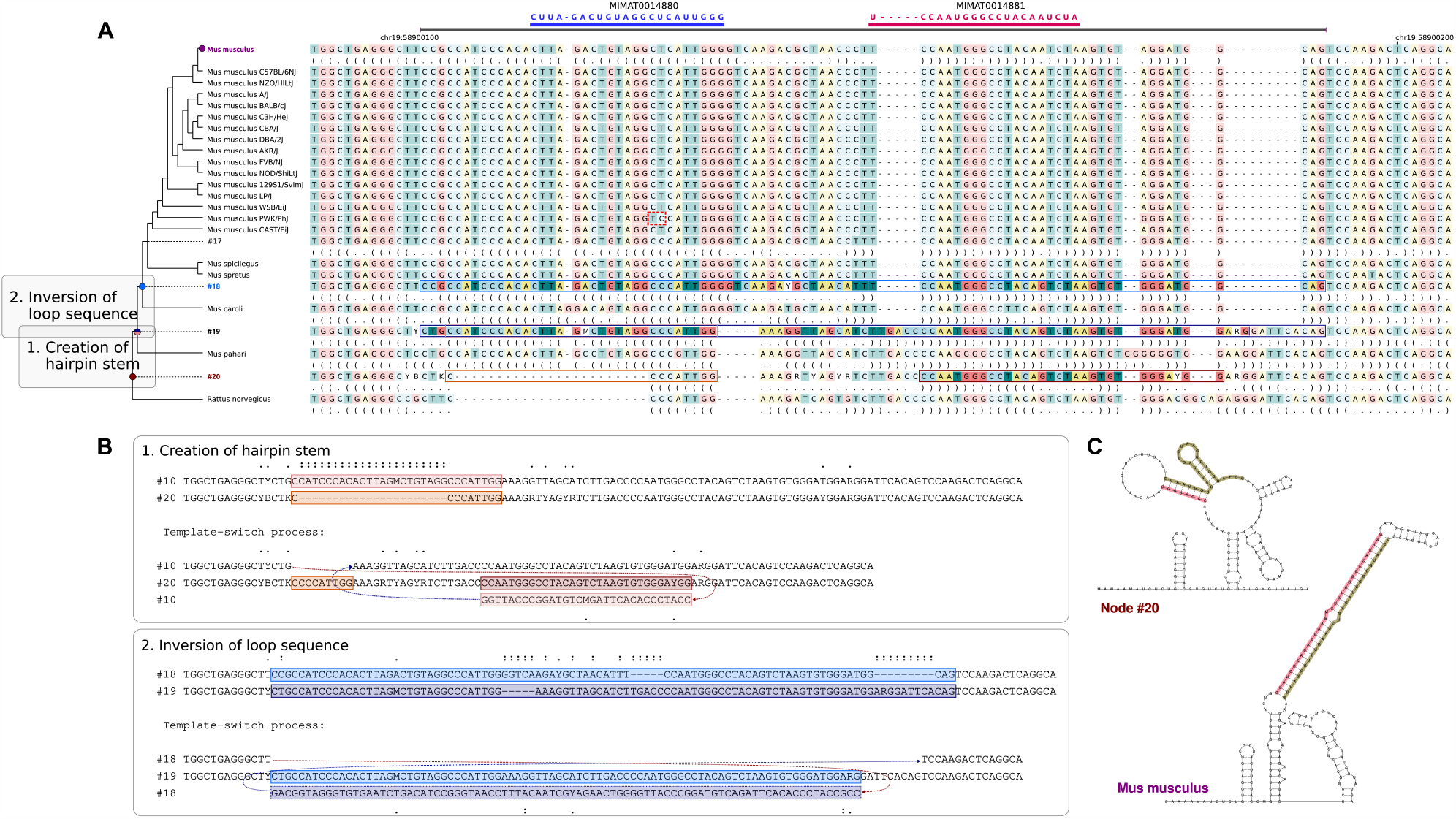
Emergence and evolution of the mmu-mir-3086 gene. (*A*) Two TSM events are inferred to have taken place in the locus among the sampled rodents. The first event in the ancestor of Murinae (between nodes #20 and #19) creates a new inverted repeat with the capacity of forming a hairpin. The second event after the split of *Mus pajari* (between nodes #19 and #18) modifies the existing hairpin by inverting the hairpin loop sequence. A two-nucleotide difference in the lab strain PWK/PhJ is indicated with a red dashed box. (*B*) The inferred template switch solutions for the two complex mutations. The sequence names (left) match the node names in the tree. (*C*) Predicted secondary structures for the sequences before and after the inferred TSM events.

### Existing promoters induce transcription of novel stem structures

Our analyses confirm that perfect stem structures can be created by single mutation events and such nascent reverse-complement repeats can further evolve into putative miRNA genes. However, the potential to form a hairpin structure is alone not sufficient for turning a TSM locus into a gene. Transcription of the hairpin sequence and its subsequent processing in the miRNA maturation pathway are both necessary for a TSM-derived sequence to function as a RISC-associated miRNA.

Transcriptional activation of the miRNA genes is associated with their genomic context. Intragenic miRNAs within protein-coding or lncRNA genes are often dependent on the host gene transcriptional activity (30) and are subsequently released from excised introns or mRNA sequences by the Drosha exonuclease (9). A subset of intragenic miRNA genes and most intergenic miRNA genes have their own promoter sequences, but intergenic miRNA may also utilise the bidirectional activity of nearby promoters or readthrough transcription of an upstream protein-coding or lncRNA gene (31–33). To differentiate between these possibilities we determined the genomic context of the TSM-derived miRNA genes (intergenic vs intragenic) and searched for genomic elements capable of providing transcription for the miRNAs. We particularly looked for SINE elements that can provide a sense promoter and LTR and LINE1 elements that can provide both sense and antisense promoters for the nearby miRNA gene.

Consistent with earlier observations of introns as hotspots for young miRNAs (29, 34), 89.2% (99/111; Table S2) of our TSM-associated miRNA loci are located in intragenic regions. A great majority of these (81/99) are in the sense strand of a host gene intron and can be expressed as a part of the transcriptional unit with a predominantly neutral effect on the host gene. Additional five miRNAs (5/99) are in the sense strand but partially overlap with an exonic region of the host gene, and of the thirteen intronic miRNA in antisense orientation to the host gene, ten (10/99) are located within 1 kb of an LTR, SINE, or LINE1 promoter. Of the twelve intergenic miRNAs (12/111), four (4/12) are located within 1 kb of a gene promoter, which could support their expression by bidirectional activity (35, 36) and the remaining eight (8/12) are within 1 kb of a transposable element (TE) promoter. In general, the nearby TEs are older than the TSM event (Table S1) but in two cases (hsamir-3678 and hsa-mir-5681) the TSM appears to predate insertions of all nearby TEs, demonstrating the independence of the appearance of a secondary structure and the gene promoter. In total, the genomic context provides a potential expression strategy for 108/111 of the TSM-associated human miRNA genes. Consistent with this, nine of the total ten TSM-generated miRNAs identified in the Murinae data are in the sense strand of a host gene intron.

The miRCarta (37) expression profiles (Fig. 2D) summarise the present RNAseq evidence for miRNA genes annotated in miRBase. Our analysis of the miRCarta profiles confirmed the expression of 110/111 of the TSM-derived miRNA genes. A closer evaluation of the read stacks revealed that at least 96 of the miRNA genes produce clearly defined miRNA-sized stacks (d*<*0.2, see Methods; Table S1, Data S1) indicating that their primary transcripts are successfully processed by the miRNA processing pathway. Fang and Bartel (8) showed that wobble pairs and a mismatched motif in the basal stem region enhance the processing of the pri-miRNAs. Consistent with this, several sequences inferred for nodes after the TSM event contain mismatches, suggesting base change mutations after the creation of a perfect inverted repeat. Further, a majority of the loci (105/111) have gained additional mutations after the initial TSM event.

Finally, we screened the miRNet (38) and RNAcentral databases for experimental validation of the candidate loci, considering miRNA expression evidence from microarray, qRT-PCR and RNA-seq analyses. Combining information from the miRNet and RNAcentral databases, we found Dicer HITS-CLIP/PAR-CLIP evidence about the gene target for 75 TSM-generated miRNAs (Table S1). For example, hsa-mir-576 gene produces a nearly optimal gene product (stack quality d=0.045; Data S1) and the RNAcentral HITS-CLIP/PAR-CLIP evidence associates it to 576 protein-coding mRNA targets.

### A promoter is a limiting factor for *de novo* miRNA genes

Our findings are in line with the suggestions of transposons being involved in the emergence of new miRNA genes (40, 41) and the reports of young miRNAs being enriched in introns of protein-coding genes (29, 34). This raises the question of whether the enrichment reflects the requirement for a pre-existing promoter or whether it results from a corresponding bias in the distribution of TSM events. To distinguish between these options, we reanalysed the candidate TSM loci found in human polymorphism data (15), shuffling the coordinates of the 3,705 TSM-explained mutation clusters across the complex parts of the genome. We found that the TSM events are strongly depleted in protein-coding genes (*p* = 2.85 × 10^*−*6^), including the introns (*p* = 0.012). Furthermore, the distances between TSM loci and the closest transposable elements do not differ from random (Fig. S4). Together, the results strongly support the enrichment of novel miRNAs in specific genomic regions reflecting the need for transcriptional expression machinery. One should note, though, that our phylogenetic approach for identifying TSM events may have favoured miRNA loci within gene regions due to their greater conservation and that we could not distinguish alternative mutation mechanisms acting on low-complexity regions. Because of the latter, we excluded all potential TSMs that overlap with repeat elements from our polymorphism data, which prevented closer analyses of the transposon loci (see Fig. S5).

To estimate the proportion of novel hairpin structures created by TSM events being converted into miRNA genes, we identified all TSMs located within genic regions in the evolutionary lineage from the primate ancestor to human. We split the human gene regions into 1 kbp blocks, fetched homologous regions from other primate genomes, aligned the sequences and scanned for TSM-like mutation patterns using a similar approach as for the miRNA loci. In total, we identified 53,666 putative TSMs of which 31,007 formed a new stem structure at the RNA level and 6,054 were within the size range of a typical miRNA gene (Table S2). Due to subtle differences in the analysis pipeline (see Methods), 57 of the originally detected 99 miRNA loci passed the quality control. Given this, our results indicate that of the human-lineage TSM loci that encode suitable hairpin structures only 0.9% (57/6,054) show sufficient evidence to be annotated as miRNA loci in miRBase (19).

## Discussion

Our analyses of microRNA (miRNA) genes annotated in miRBase (19) demonstrate that the evolutionary emergence of more than one hundred potential human miRNA genes can be traced with high confidence to DNA-replication-related template-switching mutations (TSMs) in the primate lineage. Consistent with earlier estimates (13–15), we identified abundant (>50,000) TSMs within the genic regions of the reconstructed human evolutionary lineage. Of these, more than 6,000 TSMs have the potential to produce pri-miRNA-sized hairpin structures following transcription, suggesting a potent mechanism to produce the evolutionary raw material for miRNA biogenesis (7). According to the miRCarta database, most of the TSM-generated human miRNA genes have shown some transcriptional evidence, indicating a successful passage through the miRNA processing pathway (Fig. 2D, Data S1). However, we do note that only 18 of the identified miRNA genes are included in the more stringently annotated MirGeneDB (20, 21) and are both consistently expressed and properly processed; the majority of the miRNA genes reported here lack detailed functional annotations (see Table S1), such as validated high-confidence target mRNAs or related phenotypic evidence to fulfill the most stringent definitions of miRNA genes (42). On the other hand, TSMs explain at least a quarter of the 69 miRNA genes in MirGeneDB that were inferred to have arisen in primates (25), making template switching a central mechanism in creation of novel miRNA genes.

The discrepancies between the different miRNA databases reflect both the multi-step maturation of a novel locus into a gene and the difficulties in defining what makes a real miRNA gene. Following the analogy with protein-coding protogenes (2, 43), it is possible that a subset of the TSM-associated loci annotated as miRNA genes in miRBase are proto-miRNAs and still evolving towards functionality. These young miRNAs may have acquired expression and processing characteristics but not necessarily such regulatory targets that provide a positive selection advantage and evolutionary conservation (7, 44). On the other end of the spectrum are TSM-associated miRNAs passing the stringent criteria of MirGeneDB and showing clearly documented functional significance. A noteworthy example is hsa-mir-576 (Table S1, Data S1) which has validated mRNA targets and is linked to the antiviral response threshold in primates (45).

Recent work reported the role for lncRNAs in the *de novo* emergence of 82 putative protein-coding genes in the primate lineage (4) but left the mechanism originally generating the sequences open. Similarly, whilst earlier studies have documented fast evolutionary flux for miRNA genes in several species, the mechanistic basis of *de novo* emergence of the miRNA loci has remained elusive (34, 46– 48). Our findings provide a robust model for the emergence of novel miRNA loci and their precursors. Unlike the gradual mutation process by individual single nucleotide changes (49), TSMs can generate perfect base pairing for an entire RNA hairpin in a single event. In the right context, such as in introns that are known to be enriched for evolutionarily young miRNA loci (29, 34), they would also instantly satisfy the transcriptional requirement for a functional gene. Altogether, this makes the TSM-driven process of miRNA genesis orders of magnitude faster than the *de novo* evolution of functional proteins, potentially enabling rapid rewiring of existing genetic information and quick adaptation to changing environments. Our analyses focused on past events, but many TSM variants capable of becoming miRNA genes segregate among the human populations (Fig. S6) indicating that the TSM process is active and shaping our genomes right now.

## Materials and Methods

### Data collection and preprocessing

The genetic coordinates of the human miRNA were retrieved from the miRBase (v.22.1; 19) and MirGeneDB (v. 2.1; 20, 21) databases and all unique loci were retained. The genome assemblies of 62 mammals were downloaded from the UCSC database (https://hgdownload.soe.ucsc.edu/goldenPath/archive) and Entrez (see Table S3), and their alignments were downloaded from the UCSC database (https://hgdownload.soe.ucsc.edu/goldenPath/hg38/multiz100way/maf/; Last modified 6th of May 2015). As maf-alignments consist of discontinuous homology blocks, the alignments were only used to transfer the coordinates of the human miRNA loci to other species. Of other species, we discarded ones whose concatenated homologous sequence (implied by the maf-alignment) was less than 30 bp long, showed less than 80% identity to any other sequence (defined as #identical bases/alignment length), or had incongruent orientation around the start and end coordinates. For the remaining species, the sequence homologous to the human miRNA locus with an additional 50 bp of flanking sequences was extracted from the species’ genome sequence. Three miRNAs (hsa-mir-605, hsa-mir-3688, hsa-mir-7856) are located next to a long human-specific insertion, and long enough sequences were extracted for these to ensure homology beyond the insertion gap.

The human gene coordinates were collected from the Ensemble v.105 (50) genome annotation for the human reference genome GRCh38 (http://ftp.ensembl.org/pub/release-105/gff3/homosapiens/Homosapiens.GRCh38.105.gff3.gz). The overlapping genes were merged and the remaining gene regions were split into 1000 bp long chunks with 25 bp overlap for neighboring chunks. Terminal chunks shorter than 1000 bp were merged with a previous chunk. The homologous sequences were extracted similarly to miRNAs loci with the exception that the species were limited to primates, dog, cat, pig, cow and sheep, and the minimum length of sequence implied by the maf-alignment was 100 bp.

The genetic coordinates of the mouse miRNA were retrieved from the miRBase and were transferred to the reference version GRCm39. Around each candidate miRNA locus, the Murinae EPO genome alignment (“collection-murinae”; Table S4) with 50 bp of flanking sequence and the corresponding phylogenetic tree were retrieved from Ensembl (v.110; 50) using the REST API (51).

### Ancestral sequence reconstruction and inference of TSMs

For the human miRNAs, the reference phylogenetic tree for the study species was downloaded from the UCSC database (https://hgdownload.soe.ucsc.edu/goldenPath/hg38/multiz100way/hg38.100way.nh). The sequence alignments were done in two steps. First, the sequences for a miRNA locus were aligned with PAGAN2 (52) according to the reference phylogeny. Based on this alignment, sequences were left out if 1) they were more than 1.3 times longer than the human sequence, 2) they contained ten or more Ns in a row, or 3) their sequence identity to human was *<*50% or *<*60% when using, respectively, the human and the target sequence as the reference. For the remaining sequences, the alignment and ancestral sequence inference were performed with PAGAN2 according to the same guide tree.

Phylogenetic trees were traversed with a Python script using the ete3 library package v.3.1.1 (53). Following Mönttinen et al. (18), sequences at each non-root parental node and child node were compared using the FPA tool (13) that performs a pairwise alignment using the classical forward algorithm and a novel algorithm requiring reciprocal template switches (producing an inverted fragment). Hits showing better inverted than forward alignment–with the inverted fragment ≥15 nucleotides in length, with ≥80% sequence identity and differing by ≥5 mismatches from the parent–were recorded. The secondary structures for sequences at the query node as well as its parental and children nodes were predicted using RNAfold v.2.4.14 (54) with default settings.

The identification of TSMs within gene regions was performed similarly except for quality thresholds. After the first alignment the sequence was discarded if 1) it contained more than 50 N’s in a row, 2) either end of the sequence had a gap longer than 30% of the length of the human sequence, or 3) the sequence identity to human was *<*30% when using the target sequence as the reference.

For the mouse miRNAs, each locus was realigned with PAGAN2. The resulting tree-alignment structures were similarly traversed and each parental node and child node pair was compared using the FPA tool, keeping the cases with a better inverted than a forward alignment.

### Quality control for identified TSMs

As a quality control for the identified TSM cases, the aligned sequences were compared at the mutated node (child), its ancestral form (parent) and at their sister and parent nodes; the reliability of the ancestral sequence reconstruction was separately assessed at the TSM source region (where the sequence is copied from) and at the TSM target region (where the sequence is copied to). For the primate miRNA loci, the TSM source sites were compared between 1) the mutated node and its parent; 2) its parent and sister; 3) its parent and grant-parent; 4) its grant-parent and parent’s sister; and 5) the mutated node and its children (one had to pass). The TSM target sites were compared between 1) the mutated node’s parent and sister; 2) its parent and grant-parent and 3) its grant-parent and parent’s sister. Minimum sequence identity of 80% was required between the mutated node and its parent/children, and between its parent and sister; and identity of 70% was required between its parent and grant-parent, and between its parent and sister of its parent (see Fig. S7). Uncertain bases were marked with IUPAC characters and resolved using parsimony. To confirm the inheritance of the TSM target sequence, the TSM target sequence was compared between the mutated node and its child nodes, and sequence identity of 80% was required. Finally, the TSM locus had to cover at least 95% of the human miRNA locus. With current data, the comparison between the target node’s parent and grant-parent limits the analysis to the common ancestor of primates and rodents.

Quality control for the inferred TSMs in the primate gene region data was similar with the exception that the root node was allowed to be the parent for the mutation and the comparisons of the grant-parent node were left out (Fig. S7). As repeats may sometimes look like a TSM just by accident, TSMs overlapping more than 50% of their length with simple repeat elements were excluded. Simple repeat regions were identified with dustmasker v.1.0 (55). The criteria for classifying a TSM as a new miRNA-like sequence were: 1) no equally-long hairpin was present at the same position in the parent sequence; 2) the length of the new hairpin stem was 30-55 bases; and 3) the loop of the new hairpin stem was located within the central third of the region defined by the TSM start and end sites.

The Murinae data have only one outgroup species, the rat (*Rattus norvegicus*), and similar rules for the source and target site conservation could not be applied. On the other hand, all ingroup species are from the family *Mus* and, in visual inspection, the inferences on highly similar sequences were found very good.

### Genomic enrichment of TSM events in humans

The coordinates of TSM loci segregating in human variation data with the complex genomic regions (not masked as repeats or simple sequences) were obtained from a previous study (15). Using the Ensembl v.105 (50) genome annotation (gff3) for human, the coordinates of coding genes, non-coding RNA genes, pseudogenes, coding exons, non-coding exons, and UTR3/UTR5 regions were extracted and, within each subclass, overlapping regions were merged using BEDTools v.2.26.0 (56). The genomic context of the inferred TSM loci was analyzed using a custom script. The enrichment of the events was analyzed using the Python package statistics v.3.11 and a one-sided Z-test, and the counts of inferred events in unmasked genomic regions falling into different categories were compared to the counts obtained when similar intervals were randomly shuffled across the same genomic regions.

The coordinates of transposable elements were downloaded from the UCSC database (https://hgdownload.cse.ucsc.edu/goldenpath/hg38/bigZips/hg38.fa.out.gz; Last modified 15th of January 2014). The distance between the TSM target site and the closest transposable element was computed using BEDTools. Boxplots were drawn from the results using the R package ggplot2 (57). The enrichment of TSMs in specific genomic regions was tested by comparing the original coordinates to random sets of coordinates. Shuffled sequence positions were drawn from unmasked sequence regions using BEDTools. In total, 100 replicates were generated.

### Quality of microRNA expression patterns

To assess the expression of TSM-associated miRNAs, their expression profiles, representing the number of RNAseq reads mapped on each position with zero or with one mismatch, were downloaded from the miRCarta database v.1.1 (https://mircarta.cs.uni-saarland.de; 12th of December 2022; 37). The expression pattern was visualized using the R package ggplot2 and its deviation from an optimal stack was computed with the function:

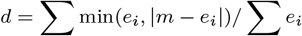

where *e*_*i*_ is the expression at position *i* and *m* = *max*(*e*_*i*_). We used the threshold *d* < 0.2 to classify the primary transcripts to be successfully processed.

### Visualization of pri-microRNA hairpin free-energies

The emergence of the stem structure was considered to be credibly explained by template switching if the solution left at most nine bases unexplained. For these cases, sequences homologous to the human pri-miRNA loci were extracted from other species and their secondary structures were predicted using RNAfold v.2.4.13 (54)with default settings. As the free-energy values vary across loci, they were normalized for visualization by dividing the inferred values by that of the greatest free energy. For the compact matrix, taxa within rodents, odd-toed ungulates, even-toed ungulates, carnivores, and afrotheres were merged and the weakest normalized value was retained. Taxa with data for fewer than ten miRNA loci were discarded. The heatmaps were generated using Python packages matplotlib v.3.5.1 and seaborn v.0.11.2.

### Identification of primate-specific miRNA loci and post-TSM mutations

The mammalian sequences homologous to the human miRNA loci were compared pairwise to the human sequence, and their identity was calculated by dividing the count of identical bases by the length of the pairwise alignment. If sequence identity higher than 90% was not found outside primates, the miRNA locus was considered primate-evolved. Post-TSM changes were similarly identified by pairwise comparison of the human sequence and the inferred sequence at the mutation node.

## Code and data availability

The Python and R scripts for data processing, analyses and visualization are available at https://github.com/helimonttinen/tsmmicrornas. The associated data and intermediate results are available at https://etsin.fairdata.fi/dataset/d3f0360c-cd36-46f5-b5ff-32398281d48d. All other data are included in the article and/or SI appendix.

## ACKNOWLEDGMENTS

We thank Brendan Battersby and Gunter Meister for the comments on an earlier version of the manuscript. This work was enabled by the Academy of Finland grant #322681 to A.L. and #341477 to M.F, and the Emil Aaltonen foundation grant to H.M. We acknowledge CSC – IT Center for Science for the computational resources.

